# Gene Deletion of the PACAP/VIP Receptor, VPAC2R, Alters Glycemic Responses During Metabolic and Psychogenic Stress in Adult Female Mice

**DOI:** 10.1101/2023.09.14.557790

**Authors:** Elena V. Kozlova, Anthony E. Bishay, Maximillian E. Denys, Bhuvaneswari D. Chinthirla, Matthew C. Valdez, Kurt A. Spurgin, Julia M. Krum, Karthik R. Basappa, Margarita C. Curras-Collazo

**Author notes:** These authors have contributed equally to this work (Anthony E. Bishay, Maximillian E. Denys). **Corresponding author:** Dr. Margarita C. Curras-Collazo, Ph.D Professor of Neuroscience Department Molecular, Cell and Systems Biology University of California, Riverside Riverside, CA 92521.

## Abstract

Pituitary adenylate cyclase-activating polypeptide (PACAP) and the homologous peptide, vasoactive intestinal peptide (VIP), participate in glucose homeostasis using insulinotropic and counterregulatory processes. These opposing actions need further characterization, as does the role of VIP receptor 2 (VPAC2R) in the regulation of glucose metabolism. In this study, we examined the participation of VPAC2R on basal glycemia, fasted glucoregulatory hormones and on glycemia responses during metabolic and psychogenic stress using gene-deleted (*Vipr2^-/-^*) female mice. The mean basal glycemia was significantly greater in *Vipr2^-/-^* in the fed state and after an 8h overnight fast as compared to wildtype (WT) mice. Insulin tolerance testing following a 5h fast (morning fast, 0.38 U/kg) indicated no effect of genotype. However, during a more intense metabolic challenge (8 h, ON fast, 0.25 U/kg), *Vipr2^-/-^* females displayed significantly impaired insulin hypoglycemia. During immobilization stress, the hyperglycemic response and plasma epinephrine levels were significantly elevated above basal in *Vipr2^-/-^*, but not WT mice, in spite of similar stress levels of plasma corticosterone. Together, these results implicate the action of upregulated counterregulatory processes influenced by enhanced sympathoexcitation. Moreover, the suppression of plasma GLP-1 levels in *Vipr2^-/-^* mice may have removed the inhibition on hepatic glucose production and the promotion of glucose disposal by GLP-1. qPCR analysis indicated deregulation of central gene markers of PACAP/VIP signaling in *Vipr2^-/-^*: upregulated medullary tyrosine hydroxylase (*Th*) and downregulated hypothalamic *Vip*. These results demonstrate a physiological role for VPAC2R in glucose metabolism, especially during insulin challenge and psychogenic stress, likely involving the participation of sympathoadrenal activity and/or metabolic hormones.

## Introduction

Vasoactive-intestinal peptide (VIP) and pituitary adenylate cyclase-activating polypeptide (PACAP) are structurally and functionally related peptide members of the secretin family, whose members include glucagon and glucagon-like peptide-1 (GLP-1) [1]. VIP and PACAP and their three class B1 G-protein-coupled receptors (GPCRs), PAC1R, VPAC1R and VPAC2R, that mediate their effects, are distributed in the CNS and peripheral tissues, and are involved in pleiotropic physiological functions. These include: immune function, pain, neuroprotection, osmoregulation, sleep/wake cycles, reproduction, stress responses and energy homeostasis [2–11]. While VPAC1R and VPAC2R are activated by both VIP and PACAP, PAC1R has a selectively high affinity for PACAP [2,3]. Due to their wide distribution these receptors, have the potential to be leveraged as therapeutic targets, however, their high structural homology has hindered the development of immunological tools and potent, selective and stable ligands, making the identification of the specific receptor subtype involved in various physiological and pathological processes difficult [12]. The use of transgenic mice lacking these peptides or receptors has offered an alternative way to study the PACAP/VIP systems [6,13]. For example, such models have allowed elucidation of the role of PAC1R in the insulinotropic response to glucose [14] and of VIP and VPAC2R in circadian rhythm [6,7,15,16].

VIP and PACAP systems have emerged as key neuropeptides regulating glucohomeostatic responses and metabolic hormones [9,17–19], with a potential implication for therapeutic intervention in diabetes and other metabolic disorders [17,20,21]. This is supported by PACAP and VIP’s role in potentiating glucose-dependent insulin secretion, leading to glucose disposal after a meal [14,22–24]. Additionally, PAC1 and VPAC2 receptors are thought to mediate the insulinotropic actions of PACAP38 and VIP on pancreatic B cells, respectively [22,23]. However, VIP and PACAP also promote an elevation in blood glucose levels. For example, exogenous administration of PACAP27 to either mice or humans increases plasma insulin without reducing plasma glucose levels [25], suggesting additional actions of PACAP on counterregulatory responses that oppose those of insulin-induced hypoglycemia. Counterregulatory actions are likely due to VPAC receptors, since administration of PACAP to PAC1R gene-deleted mice during an intravenous glucose challenge worsens glucose tolerance [14]. Indeed, PACAP and VIP application can trigger hepatic glycogenolysis and gluconeogenesis via VPAC receptors [26–29]. These counterregulatory responses produced by PACAP and VIP include direct hepatic effects but also indirect effects via secretion of the glycogenolytic hormones glucagon and epinephrine [26,27,30]. In combination, previous research reports demonstrate that VIP and PACAP and their receptors have complex actions on glucose metabolism. Importantly, VIP derivatives that selectively bind to VPAC2R are promising hypoglycemic drugs that act to promote glucose-stimulated insulin secretion and enhance glucose disposal [21,28]. However, the physiological role of VPAC2R in glucose homeostasis in vivo has not been fully elucidated and warrants further study.

Consistent with its role as a master regulator of endocrine and behavioral *stress* responses, PACAP can increase glucose levels via regulation of sympathetic nerve activity during conditions of metabolic stress [31–33]. PACAP regulates epinephrine secretion to promote hepatic glucose output and thereby can counterregulate glucose-stimulated insulin hypoglycemia, suggesting that PACAP can regulate glucose homeostasis in a biphasic manner appropriate to the physiological conditions [34]. In PACAP gene-deleted mice, insulin-induced metabolic stress (hypoglycemia) is lethal, because of inadequate adrenal epinephrine secretion [33]. Adequate epinephrine secretion requires PACAP-induced upregulation of transcripts encoding the adrenomedullary catecholamine-synthesizing enzymes, i.e., tyrosine hydroxylase (*Th*) and phenylethanolamine N-methyltransferase (*Pnmt*); this is prevented in a PACAP gene-deleted mouse model during restraint stress [35]. Several lines of evidence suggest that by modulating the release of epinephrine, glucagon and insulin from the adrenal and pancreas, respectively, PACAP, and to a lesser extent VIP, can promote sympathetic and parasympathetic function to regulate appropriate glucose responses and maintain homeostasis [24,34]. Therefore, PACAP/VIP and its receptors participate in sympathoadrenal activation in response to insulin hypoglycemic and psychogenic stress [33,36] poorly understood and understudied, especially in females [36].

Elucidating central PACAP/VIP signaling may be critical for understanding peripheral glucose homeostasis under physiological and disease states. For example, intracerebral application of VPAC2R (but not PAC1R) agonists stimulates hepatic glucose production via sympathetic innervation of the liver [37]. In the brain, PACAP is a neurotransmitter localized at stress-transducing central nuclei, including the paraventricular nucleus of hypothalamus (PVN), an area critical for the regulation of energy homeostasis and pre-autonomic control of metabolic function [36]. All 3 PACAP receptors are expressed in the PVN and in the ventrolateral medulla (RVLM), an autonomic area that receives PVN projections and is responsible for tonic and reflex control of efferent sympathetic activity, including adrenal splanchnic nerve activity [3,16,38–45]. Specifically, intra-RVLM PACAP causes PAC1/VPAC2 receptor-mediated sympathoexcitation in rats [41]. In the present study, we hypothesized that VPAC2R participates in glucose responses to metabolic and psychogenic stress and influences the expression of gene markers of PACAP/VIP signaling in central brain areas associated with its insulinotropic and sympathoadrenal functions.

In the current study, we utilized *Vipr2* gene-deleted (*Vipr2^-/-^*) females since females have augmented responses to certain stressors and are less studied in regard to PACAP/VIP signaling [46–50]. We also examined the effect of VPAC2R gene deletion on gene markers for PACAP/VIP and its receptors and associated neurotransmitters in central nuclei having glutamate (GLU)ergic pre-autonomic neurons and pre-sympathetic control neurons expressing GLU, PACAP, or catecholamines [51–53]. Compared to WT controls, *Vipr2^-/-^* female mice displayed hyperglycemia during fed and one of the fasted conditions, impaired insulin-induced hypoglycemia, and exaggerated hyperglycemia and plasma epinephrine triggered by restraint stress, implicating upregulated glucose counterregulatory processes. Gene markers for sympathoadrenal (SA) but not hypothalamo-pituitary-adrenal (HPA) axes were significantly altered in corresponding brain areas involved in VIP/PACAP signaling. Our findings are consistent with a physiological role of VPAC2 receptors in glucose homeostasis during metabolic and psychogenic stress, which likely involves the participation of sympathoadrenal activity and/or metabolic hormones.

## Materials and Methods

### Generation of Vipr2-deleted mice

Vasoactive-intestinal peptide receptor 2 gene-deleted mice (*Vipr2^-/-^*) were generated and validated as described [16]. Briefly, the *Vipr2^-/-^* mice were generated in E14/4 embryonic stem cells by replacing a 132 bp sequence containing the translation start site of the VPAC2R gene with LacZ-Neo cassette. Correct gene targeting was confirmed using RT-PCR and gene-specific primer for *Vipr2* and by radioreceptor autoradiography using the selective VPAC2R ligand Ro 25-1553. *Vipr2^-/-^* mice displayed similar gross morphology and fertility as WT littermates on a C57BL/6J background. As further validation of the mutation, we examined the expression of VPAC2R in adrenal tissue. Expression of *Vipr2* was detected in WT mice but not *Vipr2*^-/-^ mice (data not shown). *Vipr2^-/-^*mice were obtained as a gift of the last remaining colony from the Waschek Lab (University of California, Los Angeles). Twelve female *Vipr2^-/-^*and seven WT littermates were used in this study. Mice were 5.6-6.9 months of age at the beginning of the study and body weight was not significantly different between genotypes throughout the duration of the study, which lasted 10 weeks (**Supplementary Information 1**).

### Animal care and maintenance

Female *Vipr2^-/-^* and C57BL/6 wild-type (WT) control mice were maintained in accordance with the guidelines in the National Institutes of Health Guide for the Care and Use of Laboratory Animals [54]. Mice were housed in standard polycarbonate plastic cages (3-4 per cage) with corn cob bedding, unless otherwise noted. Food pellets (Laboratory Rodent Diet 5001; LabDiet, USA) and municipal tap water were provided *ad libitum* except as required during the experimental period. Temperature was maintained at 21.1–22.8°C and relative humidity at 20–70% under a 12/12 h photoperiod (lights on from 07:00–19:00 h). All experiments were approved by the IACUC on animal care and use at the University of California Riverside (AUP# 20170026 and 20200018) and the University of California Los Angeles (AUP# 93-007). Experiments were conducted between 0900-1600, unless otherwise noted. The experimental timeline is shown in **Figure 1**.

**Fig. 1.**
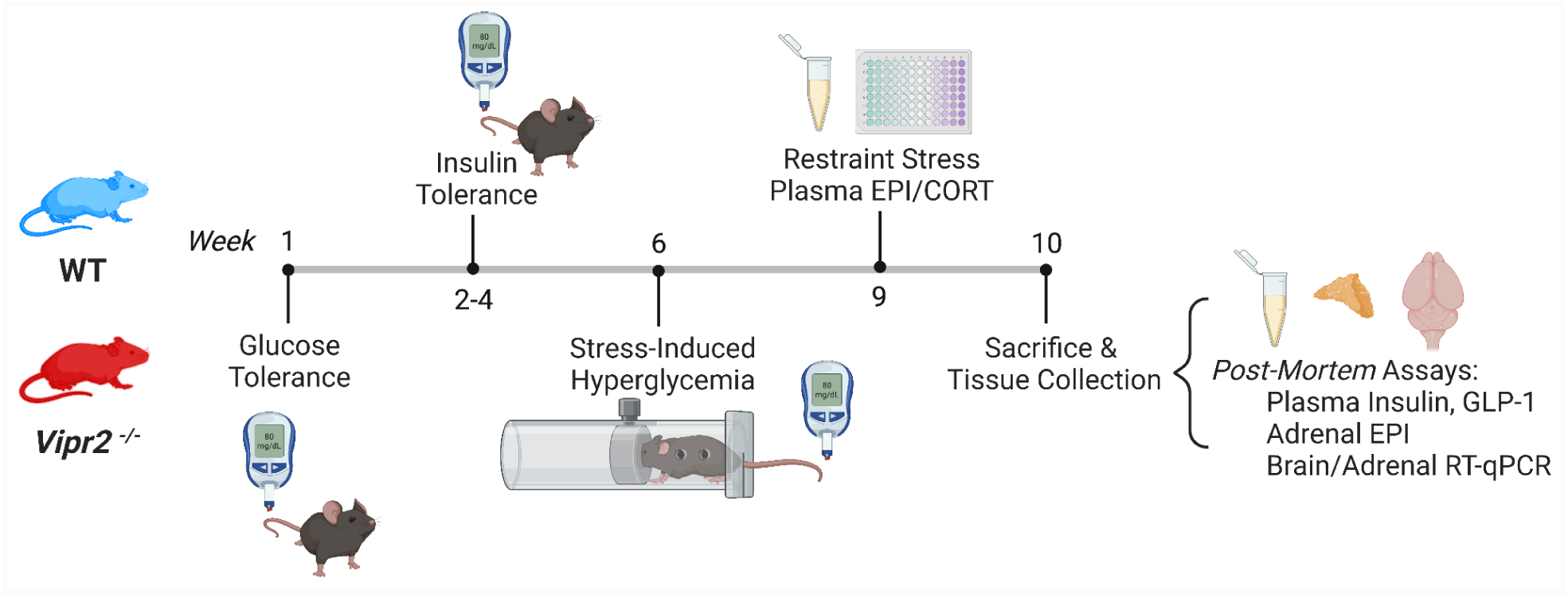
Experimental Timeline. Female *Vipr2^-/-^* mice or wildtype littermates (5.6-6.9 mo) were subjected to *in vivo* and *ex vivo* tests. At sacrifice, mice were euthanized with isoflurane and exsanguination and blood and other tissues were collected for *post mortem* analyses: ELISA, adrenal epinephrine assay and adrenal and brain RT-qPCR. CORT, corticosterone; EPI, epinephrine; GLP-1, glucagon-like peptide 1

### Glucose and insulin tolerance tests

To measure glucose tolerance (GTT), mice were fasted overnight (ON) for 11 h and glucose (2.0 g/kg b.w.) was administered by i.p. injection. Tail blood (∼1 μL) was collected at the end of the fast at t=0 (fasting blood glucose, FBG), and 15, 30, 60, and 120 min post-injection and blood glucose measured using a glucose meter (OneTouch Ultra 2, LifeScan Inc.) and corresponding test strips. One week later, an insulin tolerance test (ITT) was performed: weak challenge (morning fast, 5 h, 0.38 U/kg, i.p. of Humulin R obtained from Eli Lilly, USA or strong challenge (ON fast, 8 h, 0.25 U/kg Humulin R). Tail blood was collected at t=0, 15, 30, 45, 60, 90 and 120 min post-injection and plasma glucose measured. The area under (AUC) or above the glycemia curve (inverse AUC) obtained during GTT and ITT, respectively, was calculated over 0–120 min post injection. To determine insulin sensitivity *in vivo*, the percent blood glucose reduction rate after insulin administration, K_ITT_, was calculated over the first 30 min using the formula (0.693 x 100) × t_1/2_. Half-life (t_1/2_) was calculated from the slope of the blood glucose concentration response during 0–30 min post insulin injection, when plasma glucose concentration declines linearly [55].

### Stress-induced hyperglycemia

Two weeks after ITT, mice were subjected to immobilization stress for 90 min by restraint in devices made from 50 ml clear polystyrene conical tubes, which were perforated with numerous air holes for ventilation in a quiet, well-ventilated room at 22.7°C. Tail blood was sampled while mice were restrained at t=0, 15, 30, 60, 90 min after the start of restraint and following a 30-min rest (at t=120 min) [56], [57]. The area under the stress glycemia curve was calculated over 0–120 min.

### Measurement of plasma glucagon-like peptide and insulin

Plasma collected via cardiac puncture at necropsy (fasted state) was analyzed using commercial ELISA kits according to manufacturer’s instructions for insulin (Cat. #80-INSMSU-E01, ALPCO) and total glucagon-like peptide-1 (GLP-1) in both the 7-36 and 9-36 forms (Cat #43-GPTHU-E01 ALPCO). For insulin, the colorimetric reaction product was read as optical density at 450 nm on a plate reader (SpectraMax 190, Molecular Devices, San Jose, CA USA). The ALPCO insulin ELISA had a sensitivity of 0.019 ng/ml in a standard range of 0.025–1.25 ng/ml. GLP-1 was detected by indirect sandwich amide chemiluminescence ELISA using a luminescence plate reader (GloMax Promega, USA). This total GLP-1 assay detects GLP-1 (7-36) 100% and GLP-1 (9-36) 100% but not GLP-1 (9-37) (< 0.1%), GLP-1 (7-37) (< 0.1%), GLP-1 (1-36) (< 0.1%), GLP-2 (< 0.1%), nor glucagon (< 0.1%). This assay had an analytical sensitivity of 0.6 pmol/L in a standard range of 2.1-54 pmol/L. Three samples from *Vipr2^-/-^* displayed GLP-1 concentrations below the method detection limit (MDL). For these samples random numbers were generated around this minimum (determined as 0.249518 pmol/l) using the standard deviation of the entire group (0.033250).

### Measurement of plasma corticosterone and epinephrine

Three weeks after stress-induced hyperglycemia, tail blood (30-80 μL) was obtained immediately following a 1-h restraint session and collected into heparinized hematocrit tubes and stored on ice. Plasma epinephrine was measured using a commercial competitive enzyme-linked immunosorbent assay (ELISA) kit (Eagle Assays, ADU39-K01). After blood collection, plasma was separated via refrigerated centrifugation, stabilized with the Sample Stabilizer reagent provided in the kit and stored at -80°C until further use (within 10-14 d). Within 10-14 d of plasma collection, 8-30 μl of plasma was removed from each sample and used for the epinephrine assay. This assay is specifically designed for mouse/rat plasma and has a sensitivity of 1.6 pg/ml in a standard range of 0.15-25 ng/ml.

The same plasma samples were diluted 10-fold and corticosterone (CORT) was measured using a commercial competitive ELISA kit (Abcam, AB108821). The assay had a sensitivity of 0.30 ng/ml in a standard range of 0.39 ng/ml to 100 ng/ml. The unknown plasma epinephrine and CORT concentrations of test samples were determined by plotting their absorbance values and interpolating the concentration using a 4-parameter-logarithmic standard curve.

### Trihydroxy indole (THI) catecholamine assay

Epinephrine content in adrenal glands was measured using a modification of the trihydroxyindole method as described [58]. Using this method of catecholamine oxidation at 0°C, Kelner and colleagues obtained values for epinephrine content in bovine chromaffin cell lysates that were nearly identical to those measured using HPLC/electrochemical determination [59]. Briefly, adrenal tissue homogenates were centrifuged at 15,000 × *g* at 0°C for 15 min. Sample supernatant was incubated with 10% acetic acid. Then 0.25% K_2_Fe(CN)_6_ was added to each sample and the mixture was incubated at 0°C for 20 min. The oxidation reaction was stopped by the addition of NaOH solution containing alkaline ascorbate. Fluorescence emission was determined at 520 nm using a fluorescence plate reader (GloMax, Promega, USA). Each sample yielded mean fluorescence intensity units that were converted into epinephrine concentration expressed as μg/g adrenal wet weight using calibration standards and polynomial curve fitting.

### Brain RNA extraction

At sacrifice, using isoflurane anesthesia and cervical dislocation, whole brains and adrenal glands were rapidly dissected and snap-frozen in 2-methylbutane over dry ice. The unilateral hypothalamus and medulla oblongata were dissected by a trained neuroanatomist and immediately homogenized in TRIzol Reagent (Ambion by Life Technologies, USA) using a hand-held homogenizer (Biospec Bio-vortexer). Total RNA was prepared via a guanidinium thiocyanate–phenol–chloroform extraction followed by manufacturer’s instructions, including DNase 1 treatment (Monarch Total RNA Miniprep kit, #T2010, New England Biolabs, USA and QIAGEN RNeasyMini kit, #74104). Purity and quantity of RNA were assessed by determining the optical density (OD) photometrically using 260/280 nm and 260/230 nm ratios (NanoDrop ND-2000, Thermo-Fisher Scientific Inc., Waltham, MA, USA). Mean RNA yield was 230 ng/μL (range: 94–454 ng/ul). RNA integrity was assessed using automated electrophoretic analysis (2100 Bioanalyzer, Agilent Technologies Inc. Santa Clara, CA, USA) and yielded values with a mean of 8.5 (range 7.7–9) (**Supplementary Information 2**).

### Quantitative polymerase chain reaction (RT-qPCR) analysis

RT-qPCR was used to quantitate mRNA transcripts for the following genes: *Th*, *Crh*, *Pacap*, *Pac1r*, *Vip*, *Vglut2*. DNA oligonucleotide PCR primers were custom-or pre-designed (Integrated DNA Technologies, Coralville, IA, USA). Custom primers were designed to meet several criteria using NCBI Primer Blast followed by running of the complementary DNA (cDNA) synthesis product made using RT-PCR (iScript kit, Cat no. 1708890, Biorad, USA) on an electrophoresis gel. Only primers that gave single-band amplicons in the presence of RT and that matched the base length of the predicted target were selected. In addition, primers selected yielded 87.7-104.2% efficiency on RT-qPCR (**Table 1**). A temperature gradient to capture ideal annealing temperatures for custom designed primer pairs. The primer concentration ranged from 200-500 nM. RT-qPCR was performed on RNA (10 ng) samples, run in triplicate, on a CFX Connect or CFX96 (Bio-Rad, USA) thermocycler with the Luna Universal one-step qPCR Master Mix (E3005; New England Biolabs, Ispwich, MA, USA). Amplification reactions for genes of interest were performed in 50 cycles of the following cycling protocol: reverse transcription 55°C/10 min; initial denaturation 95°C/1 min; per cycle 95°C/10 s denaturation, 60°C or 55 °C/30 s extension; 65-95°C in 0.5°C, 5s increments melt curve analysis. To rule out extraneous nucleic acid contamination and primer dimer formation, no template controls were included in each experiment. Additionally, to rule out presence of genomic DNA (gDNA), negative RT controls, which contained the complete RNA synthesis reaction components without the addition of the enzyme reverse transcriptase (RT) were included. qPCR experiments were performed in adherence to MIQE guidelines [60] (**Supplementary Information 2**).

**Table 1.**
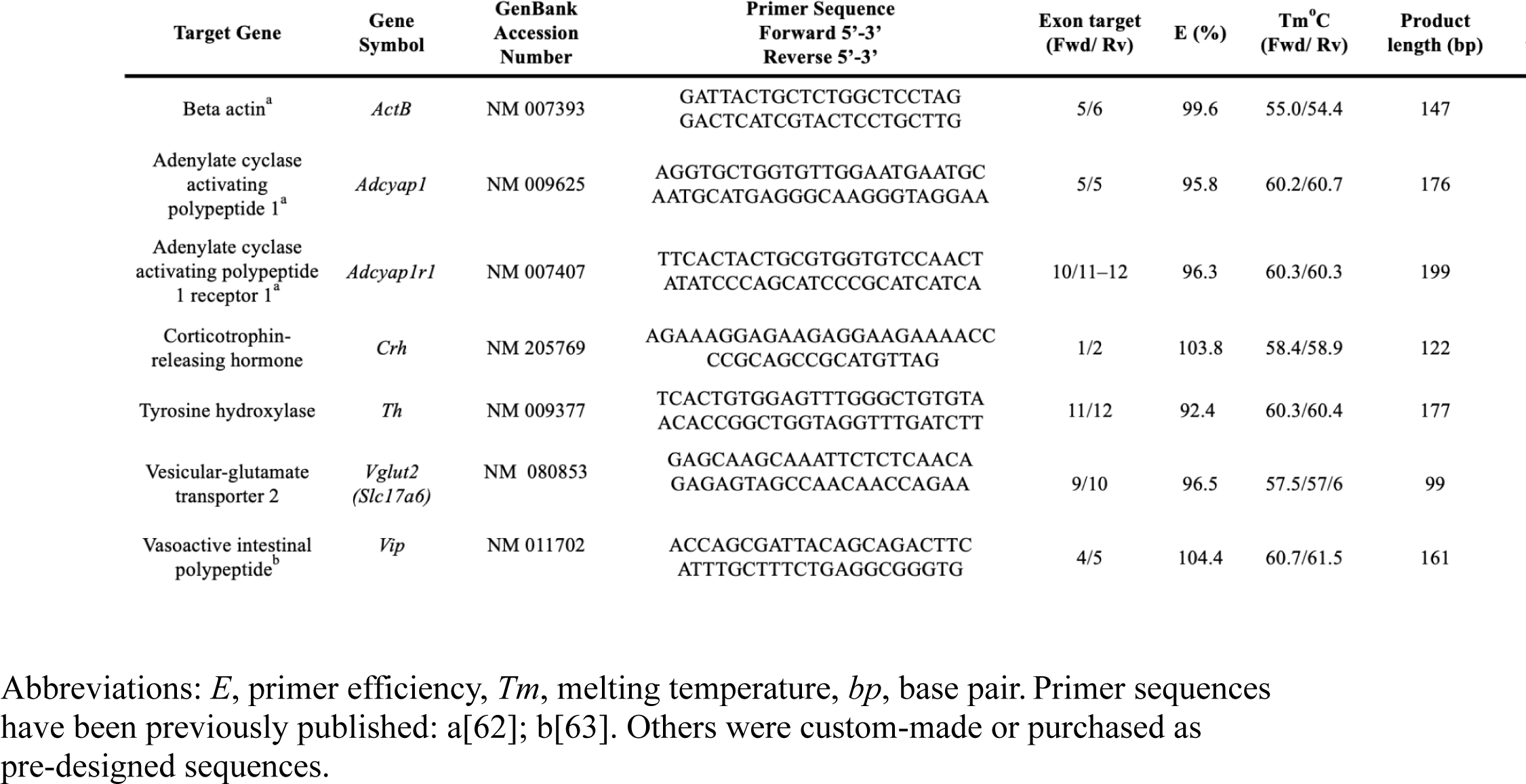
RT-PCR primer sequences, efficiencies and PCR products.

Gene expression was measured relative to the reference gene, *ActB*, and differential gene expression was determined compared to WT using the Pfaffl method [61]. *Vipr2* gene deletion did not affect the stability of the *ActB* reference gene. Mean (± s.e.m.) cycle quantification (Cq) values for medulla were 18.81 ± 0.34 and 18.79 ± 0.22 for WT and *Vipr2^-/-^*mice, respectively. Cq values for hypothalamus were 18.33 ± 0.20 and 18.38 ± 0.15, respectively. Cq values for adrenal were 17.90 ± 0.27 and 17.93 ± 0.39, respectively. Since *Vipr2* gene deletion produced an apparent reduction in adrenal weight, data are presented as both relative gene expression and copy number/ng total RNA per mg of fresh adrenal tissue obtained at sacrifice. To calculate copy number, for each sample, the corresponding ng dsDNA was determined for the mean sample Cq using the GOI standard curve. The value was inserted into the following equation that uses Avogadro’s number, length of amplicon (in bp) and average mass of 1 bp dsDNA (660 ng) to calculate the mean absolute transcripts value (copy number):

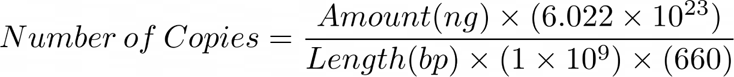

### Statistical analysis

All statistical analyses were conducted using GraphPad Prism (GraphPad Software version 9.5.1 for Mac OS X, San Diego, CA, USA). Bar graphs include background columns indicating group mean. Error bars in bar graphs and scatter plots represent s.e.m. Sample sizes are indicated in the figure captions. Differences were deemed statistically significant at *p*<0.05. Student’s *t-*test was used to examine the effects of VPAC2R gene deletion. A non-parametric t-test (Mann Whitney U test) was used when normality assumptions of at least one treatment group were not met, as assessed by the Shapiro Wilk or Kolmogorov-Smirnov tests for normality. Welch’s t-test was used when standard deviations were significantly different. The effect of *stress* or *time* and *genotype* and their interaction were determined using a two-way analysis of variance (ANOVA) or a mixed-effects model with or without repeated measures (RM). Where the F ratio was significant, *post-hoc* comparisons were assessed using the Holm-Sidak’s or Sidak’s tests for multiple comparisons. For PCR analysis, within each brain region, the False Discovery Rate (FDR) of 5% was used to correct for type I error as a result of multiple comparisons. In these cases, adjusted *p* values were calculated using the Benjamini-Hochberg test. Pearson’s correlation and linear regression were used to quantify linear relationship between variables. Pearson’s coefficient and coefficient of determination were considered significant at *p* ≤ 0.05.

## Results

### Vipr2 gene deletion produced fasting hyperglycemia

Pre-diabetes is characterized by abnormally high fasting blood glucose levels. In order to examine the effect of *Vipr2* gene deletion on glucose metabolism, blood glucose levels were examined after 5 (AM), 8 (ON) and 11 (ON) hours of fasting as well as in the *ad libitum* fed state. **Table 2** illustrates that mean glycemia values for the *ad libitum* fed state were significantly greater in *Vipr2^-/-^* than WT mice (Student’s t-test: *t*_(17)_ =1.78, *p* = 0.047). This was also the case after an 8 h ON fast. *Vipr2^-/-^* displayed reduced glycemia after a 5 h AM fast but not after an 8 h ON fast when compared to ad *libitum fed* state. In contrast, WT values remained similar to their corresponding baseline. A repeated measures (RM) two-way mixed effects ANOVA revealed a statistically significant effect of *treatment* (*F*_(3,51)_ =5.24; *p*=0.003) and *genotype* (*F*_(1,17)_ = 3.52; *p*=0.08) but not interaction (t*reatment × genotype F*_(3,51_=1.65, ns). A Holm-Sidak’s *post hoc* test for multiple comparisons revealed a significantly increased hyperglycemia in *Vipr2^-/-^* vs WT at 8 h ON (*p*<0.027) and within *Vipr2^-/-^* baseline vs 5 h ON (*p*<0.004) and 5 h AM vs 8 h ON (*p*<0.01). No significant effects were discovered for WT.

**Table 2.** Vipr2 gene deletion produced blood glucose alterations during fed and fasting conditions. Fasting blood glucose measurements (mg/dL) were measured using tail blood drawn under various fasting/fed conditions: *Ad libitum* food (baseline), 5 h morning (AM; during inactive period), 8 h and 11 h overnight (ON; during active period). Baseline values were obtained prior to fast onset. *significantly different vs WT (*p*<.05). ^significantly different vs corresponding baseline (^^*p*<.01). ^@^significantly different vs corresponding 5 h AM, (^@@^*p*<.01), *n*, 7-12/group. Values are mean ± s.e.m.

|  | *Ad libitum* food | 5 h AM | 8 h ON | 11 h ON |
| --- | --- | --- | --- | --- |
| WT | 176.6 ± 7.8 | 148.1 ± 8.5 | 147.0 ± 11.6 | 175.9 ± 16.7 |
| *Vipr2*<sup>-/-</sup> | 193.1 ± 6.5* | 151.8 ± 5.1^^ | 189.0 ± 10.1*@@ | 184.1 ± 13.0 |

### Vipr2 gene deletion did not affect glucose tolerance

To more fully investigate glucose metabolism in *Vipr2* gene-deleted female mice, blood glucose was measured during IPGTT. Glucose injection raised endogenous blood glucose levels predictably with no genotype effect (two-way RM mixed effects ANOVA: *time* (*F*_(4, 66)_ =132.3, *p* < 0.0001); *genotype* (*F*_(1,17)_ = 0.89, ns); *time × genotype* (*F*_(4,66)_ = 0.74, ns). For both groups, glycemia was significantly greater at 15-60 min post-injection relative to baseline and returned to normal by 120 min (**Fig. 2A**). There were no group differences in magnitude or duration of glycemia represented as the area under the glucose curve, AUC_IPGTTglucose_ (Welch’s t-test: *t*_(14.6)_ = 0.15, *p* = 0.88) (**Fig. 2B**). Interestingly, some individual *Vipr2^-/-^*mice showed peak glycemia at 30 min, whereas all of the WT mice had peak responses sooner post injection at 15 min (**Fig. 2C**, Mann-Whitney U=24.50, n_1_=15, n_2_=15, adjusted *p* =0.06).

**Fig. 2.**
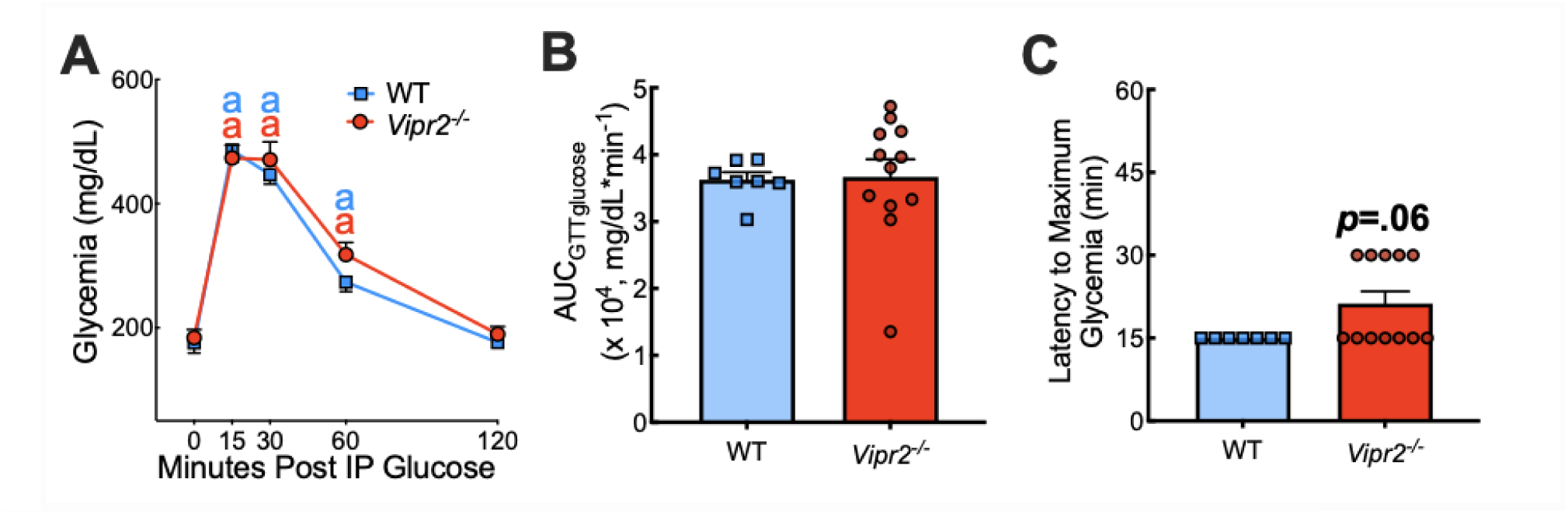
Vipr2 gene-deletion produced delay in peak glycemia after glucose challenge. **(A-C)** Mice were fasted for 11 h overnight and tail blood was sampled for glucose before (*t* = 0 min) and after (*t* = 15, 30, 60 and 120 min) i.p. injection of 2.0 g/kg glucose (IPGTT). (**A**) Absolute blood glucose concentrations taken during IPGTT in *Vipr2^-/-^* and WT (IPGTT_glucose_). (**B**) Corresponding mean values for the integrated area under the IPGTT glucose curve (AUC_IPGTTglucose_). (**C**) Latency to maximum glycemia during IPGTT. ^a^significantly different from corresponding baseline (^a^*p*<0.01-0.0001). *n*, 7-12/group

### Vipr2 gene deletion impaired insulin-induced hypoglycemia

Next, we examined the glycemia response to exogenous insulin during ITT experiments. When a less stringent insulin challenge was used, i.e., 5 h morning fast with 0.38 U/kg insulin, mean blood glucose levels dropped and then partially recovered post injection. RM two-way ANOVA indicated main effects of time and genotype (*time* (*F*_(2.14,32.05)_ = 25.8, *p* <0.001); *genotype* (*F*_(1,15)_ =0.008, ns; *time × genotype* (*F*_(6,90)_ =0.65, ns). A Holm-Sidak *post-hoc* test indicated significant drop in glycemia over time post injection. There were no differences between *Vipr2^-/-^*and WT mice in K_ITT_ nor inverse AUC_ITTglucose_ (**Fig. 3A-C**).

**Fig. 3.**
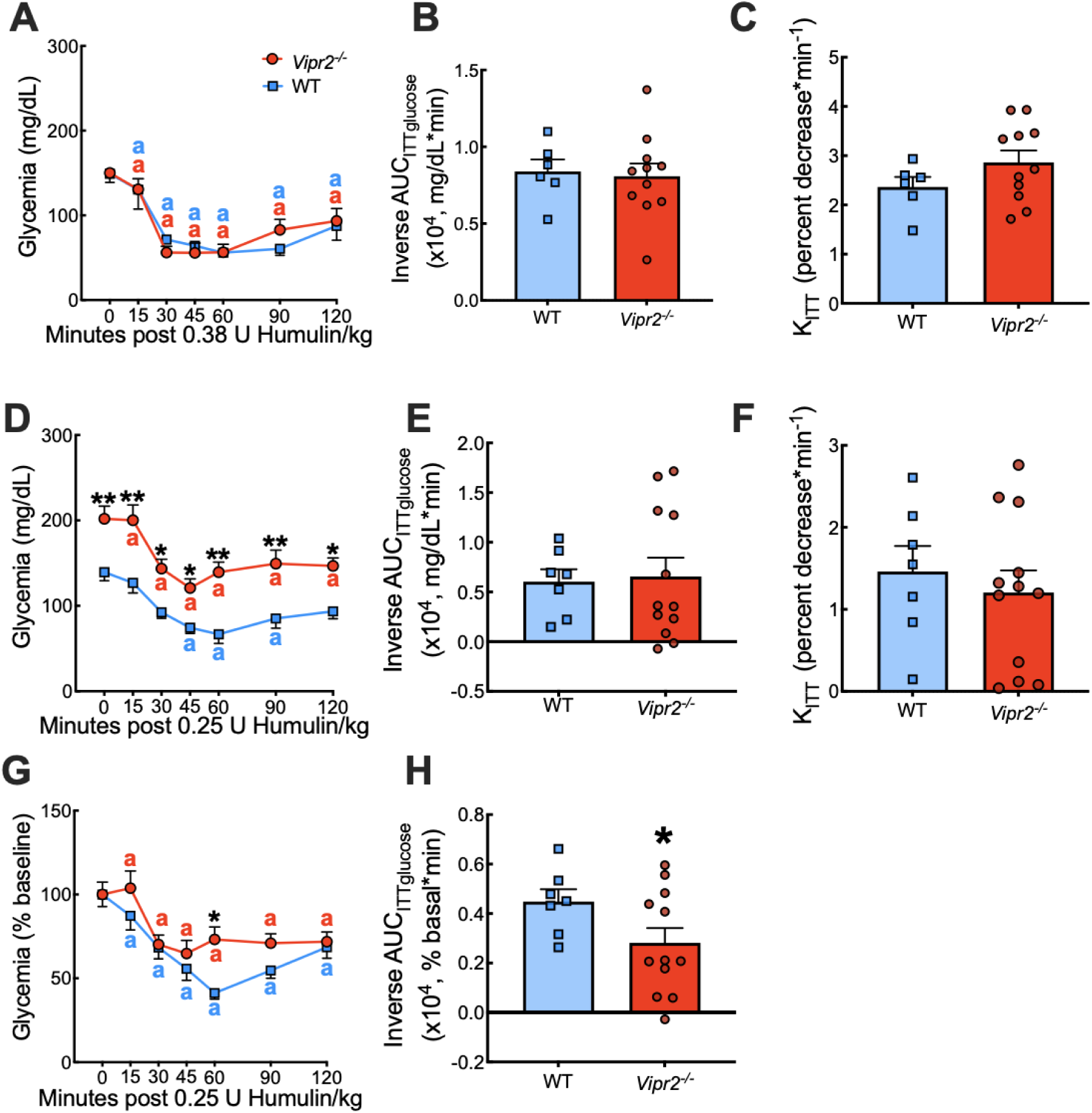
*Vipr2* gene-deletion produced a differential response to insulin challenge. Absolute blood glucose concentrations before and at *t*=15, 30, 45, 60, 90 and 120 min post i.p. insulin for female *Vipr2^-/-^* and WT mice were recorded after weak hypoglycemic stress **(A-C)** (5 h morning fast, 0.38 U/kg Humulin) and strong hypoglycemic stress (8 h ON fast, 0.25 U/kg Humulin) **(D-H)**. (**A**) Absolute blood glucose response to hypoglycemic stress (5 h morning fast, 0.38 U/kg Humulin). (**B**) Glycemia was analyzed by an inverse integrated area under the ITT glucose curve (AUC_ITTglucose_). (**C**) The corresponding rate constant for percent glucose reduction (K_ITT_) was calculated over the initial slope of the ITT glucose response curve from 0-30 min post-injection. (**D**) Absolute blood glucose concentrations were taken over the post-injection time course after strong stimulation. (**E**) Corresponding mean values for the inverse integrated area under the ITT glucose curve (AUC_ITTglucose_) are plotted. (**F**) The early effects of insulin, represented as K_ITTinsulin_, were measured over the first 30 min post-injection. (**G**) Glucose values taken during ITT are plotted versus time as a percentage of the individual basal level. (**H**) The corresponding inverse integrated area (AUC) under the percent basal glucose curve (AUC_ITTglucose_). *significantly different vs WT, \**p*<0.05, \*\**p*<0.01. ^a^significantly different from corresponding baseline (^a^*p*<0.05-0.0001). *n*, 6-12/group

Following 8 h ON fast and 0.25 U/kg insulin, there was a reduction in glycemia over time in both groups with subsequent return to near baseline levels in WT only (**Fig. 3D**). RM two-way ANOVA indicated main effects of time and genotype (*time* (*F*_(6,102)_ =12.72, *p* <0.0001); *genotype* (*F*_(1,17)_ =24.95, *p*=0.001; *time × genotype* (*F*_(6,102)_ =0.43, ns). A Holm-Sidak *post-hoc* test indicated that *Vipr2^-/-^* displayed elevated glycemia at all post-injection timepoints vs WT (*p<*0.05-0.01). Compared to baseline glycemia levels were lower at 60 min for both groups (p<0.05-0.0001) and recovered in WT but not *Vipr2^-/-^* by 120 min. However, the inverse AUC_ITTglucose_ and K_ITT_ indicate no group differences (**Fig. 3E,F**). Since these results were likely confounded by the elevated fasting blood glucose in *Vipr2^-/-^* mice (**Table 2**), we expressed glycemia as percent baseline in **Fig. 3G** (Two-way ANOVA: *time* (*F*_(6,102)_ =13.02, *p* =0.0001); *genotype* (*F*_(1,17)_ = 3.21, ns); *time × genotype* (*F*_(6,102)_ =1.41, ns). In this case, Mean glycemia values for *Vipr2^-/-^* were significantly elevated vs WT, indicating an impaired insulin hypoglycemia at 60 min post-injection (Sidak’s *post hoc*, *p*<0.05). The mean inverse AUC_ITTglucose_ (**Fig. 3H**) revealed a markedly lower glucose response to insulin for *Vipr2^-/-^* relative to WT, (Student’s t-test: *t*_(17)_ =1.90, *p* =0.04).

### Vipr2 gene deletion elevated plasma insulin and lowered GLP-1

Having observed elevated fasting blood glucose and inadequate hypoglycemia during insulin challenge, we tested whether *Vipr2^-/-^* mice displayed changes in plasma insulin and GLP-1 using ELISA. Plasma insulin levels were significantly greater in *Vipr2^-/-^* vs WT (Welch’s t-test: *t*_(7.03)_ =2.18, *p* =0.03) (**Fig. 4A**). Plasma GLP-1 levels were significantly lower in *Vipr2^-/-^* vs WT (Student’s t-test: *t*_(11)_ =3.17, *p* =0.009) (**Fig. 4B**).

**Fig. 4.**
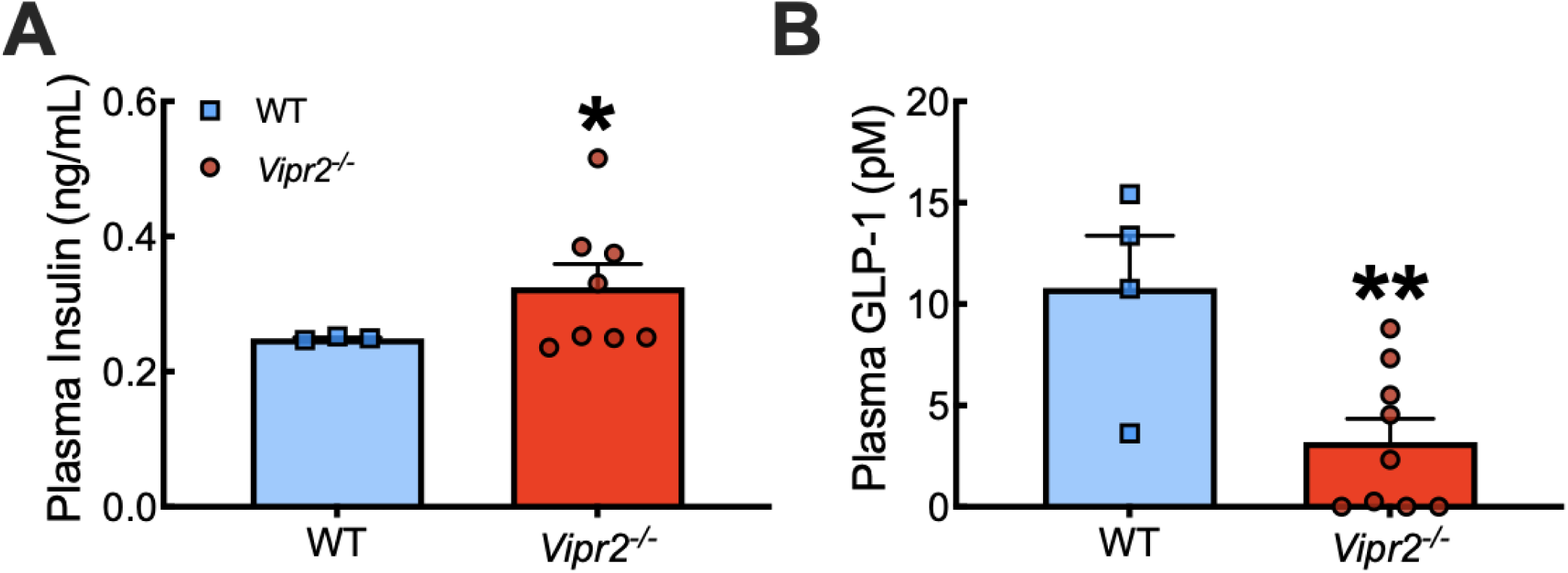
*Vipr2* gene deletion elevated plasma insulin and lowered GLP-1. Blood collected at sacrifice (fasted state) obtained from *Vipr2^-/-^* and WT female mice was assayed for plasma levels of insulin (**A**) and glucagon-like peptide 1 (GLP-1) (**B**) using antigen specific ELISAs. *Vipr2^-/-^*displayed elevated insulin and reduced GLP-1. *significantly different vs WT, \**p*<.05, \*\**p*<.01. *n*, 3-9/group

Correlation and regression analysis for stress glycemia (AUC) vs plasma GLP-1 yielded an apparent negative correlation in WT: r=-0.83, *p*=0.04, R^2^=0.70, *p* =0.08 for WT and *r*=-0.41, *p*=0.13, R^2^=0.17, *p*=0.27 for *Vipr2^-/-^* (**Fig. 8D**). These results may indicate that high stress glycemia values are associated with low GLP-1. Because this did not occur in *Vipr2* gene-deleted mice, gene deletion may be interfering with GLP-1 glucoregulatory actions. Additionally, GLP-1 was not correlated to impaired insulin hypoglycemia nor to plasma insulin levels suggesting unrelated regulation of these parameters in our gene-deleted mice (**Supplementary Information 1**).

### Vipr2 gene deletion produced exaggerated hyperglycemia during restraint stress

To examine the effects of *Vipr2* gene deletion on glucose metabolism under sympathetic overactivation, glycemia was measured during 90 min of acute restraint stress and after a 30 min post-stress period (GlucoseRes). Female WT C57BL/6J mice normally display an elevated plasma glucose level of ∼240 mg/dL in response to immobilization as shown in our study [64]. In our study this stress paradigm elevated blood glucose levels within 15-30 min for both groups (**Fig. 5**). Mean (± s.e.m.) peak glycemia was 235±18.6 and 297±13.6 for WT and *Vipr2*^-/-^ female mice, respectively. RM two-way ANOVA yielded a t*ime* effect (*F*_(3.78_, _64.19)_ = 19.59, *p* = 0.0001) and *genotype* effect (*F*_(1,17)_=10.89, *p* = 0.004) but not *time × genotype* interaction (*F*_(5,85)_=1.80, ns). Holm-Sidak’s *post hoc* comparisons indicated that, relative to WT, *Vipr2^-/-^* displayed significantly elevated glycemia at 15 min restraint (*p*<0.01) (**Fig. 5A**). The glycemic response for *Vipr2^-/-^* continued to be greater than baseline at all subsequent time points during restraint (**Fig. 5A,** *p*<0.01-*p*<0.001). *Vipr2^-/-^* glycemia values normalized only after 30 min of rest following restraint. In contrast, WT restraint glycemia was elevated over baseline at only one timepoint (30 min restraint, *p*<0.05). The exaggerated stress-induced glycemia observed in *Vipr2^-/-^* mice was reflected in the area under the restraint glucose curve AUC_GlucoseRes_ (Student’s t-test: *t* _(17)_=3.05, *p*=0.007) (**Fig. 5B**).

**Fig 5.**
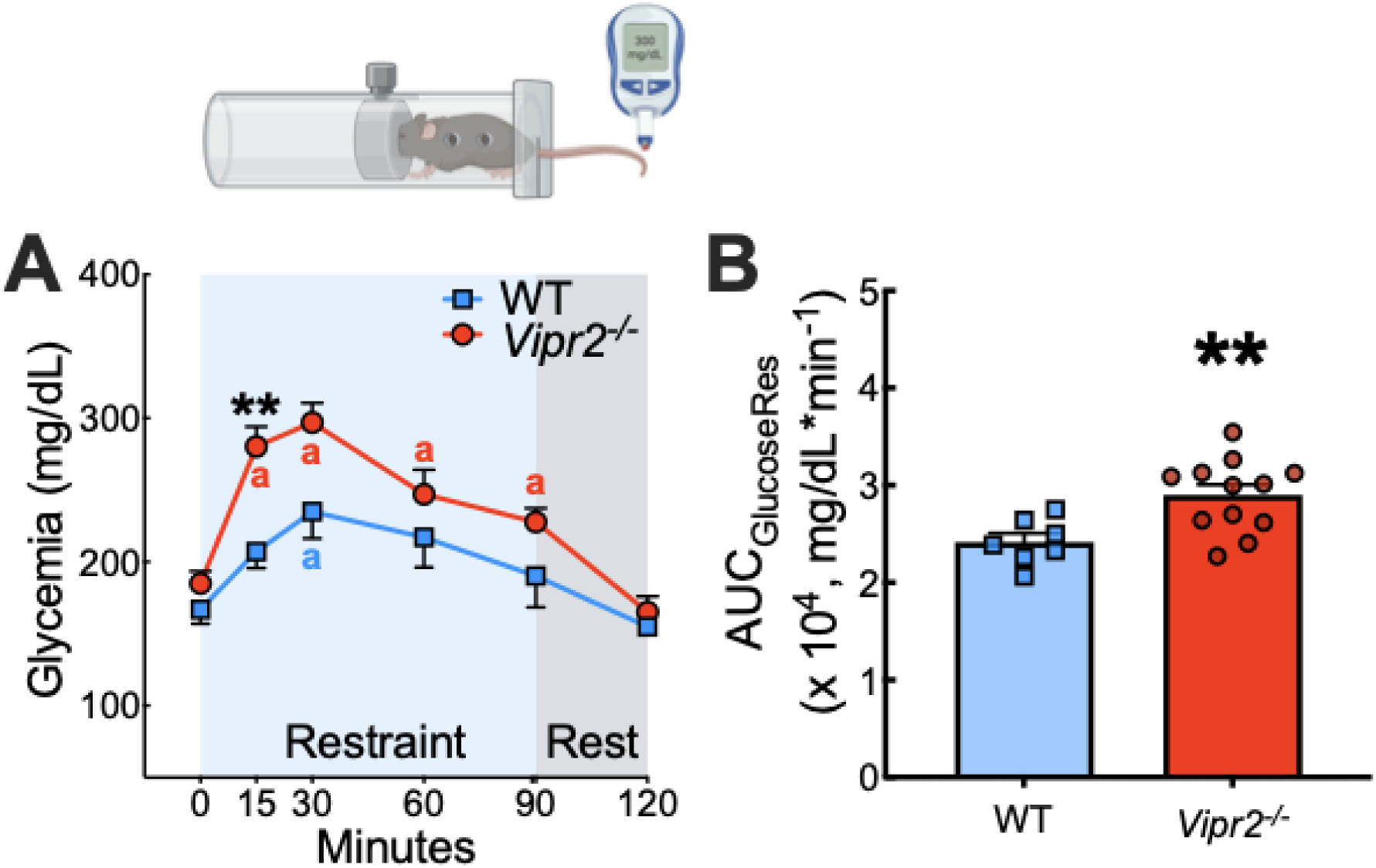
Acute restraint stress produced exaggerated hyperglycemia in *Vipr2^-/-^* mice. **(A-E)** Tail blood glucose in *ad libitum* fed mice was sampled before (*t*=0) and during (*t*=15, 30, 60 and 90 min) restraint stress (blue shading), and after 30 min of rest (grey shading). (**A**) Mean absolute blood glucose concentrations taken during the restraint stress experiment (GTT_ResGlucose_). (**B**) Corresponding mean values for the integrated area under the restraint glucose curve (AUC_ResGlucose_). *significantly different vs WT, \*\**p*<0.01. ^a^significantly different from corresponding baseline (^a^*p*<0.01-0.0001). *n*, 7-12/group

Correlation and regression analysis showed an apparent *positive* relationship found between insulin glycemia (inverse AUC) vs stress glycemia (AUC) for mutant mice only: *r*=0.52, *p*=0.04, *R^2^*=0.27, *p*=0.08 for *Vipr2^-/-^* and r=-0.29, *p*=0.26, R^2^=0.09, *p*=0.52 for WT (**Fig. 8A**). Figure 8B shows an apparent negative association between insulin glycemia (nadir) vs stress glycemia (peak) in mutants but not WT: *r*=-0.51, *p*<0.05, R^2^=0.26, *p*=0.09 for *Vipr2^-/-^* and r=-0.39, *p*=0.19, R^2^=0.15, *p*=0.39 for WT.

Correlation and regression analysis between stress glycemia (peak) vs hypothalamic *Vip* gene expression yielded a linear relationship with good fit (r=-0.71, *p*=0.04) and an apparent negative association (R^2^=0.49, *p*=0.08) in WT (**Fig. 8E**). For mutants there the linear fit was only apparent (*r*= 0.45, *p*=0.07) as was the positive association (R^2^=0.20, *p*=0.14).

### Vipr2 gene-deletion produced exaggerated stress-induced plasma epinephrine but not corticosterone

Given that *Vipr2^-/-^* mice displayed exaggerated stress-induced hyperglycemia we tested for group differences in plasma levels of stress hormones after 1 h of restraint stress. At this time point, mean plasma epinephrine was significantly elevated relative to baseline in *Vipr2^-/-^* but not WT mice. A two-way ANOVA revealed a significant effect of *stress* (*F*_(1,15)_ =9.54, *p*=0.008), *genotype* (*F*_(1,15)_=5.8, *p*=0.03) but not *stress* x *genotype* interaction (*F*_(1,15)_ =1.42, ns). A Holm-Sidak’s *post-hoc* test for multiple comparisons showed a mean elevation in plasma epinephrine in stress vs sham that was significant for *Vipr2^-/-^* (*p*=0.006) and apparent for WT (ns) (**Fig. 6A**). Mean epinephrine values during restraint were significantly elevated in *Vipr2^-/-^* compared to WT (*p*=0.04). These results suggest that *Vipr2* gene deletion exaggerates sympathoadrenal epinephrine secretion after acute psychogenic stress. To determine if *Vipr2* gene deletion affects levels of adrenal epinephrine, we compared group mean values at sacrifice. Adrenal epinephrine content was significantly lower in *Vipr2^-/-^* compared to WT mice (Mann-Whitney U=8, n_1_=624.3, n_2_=170.9, *p* =0.0013) (**Fig. 6C**). This occurred without a reduction in adrenal weight compared to WT (Student’s t-test: *t*_(17)_ =1.92, *p*=0.07) (**Fig. 6D**). Correlation analysis showed no significant relationship between stress hyperglycemia and plasma epinephrine in either group (**Supplementary Information 1**). However, an apparent negative relationship (r=-0.71, *p*=0.06 for *Vipr2^-/-^*) was found between plasma and adrenal epinephrine in the mutants only (**Fig. 8C**).

**Fig 6.**
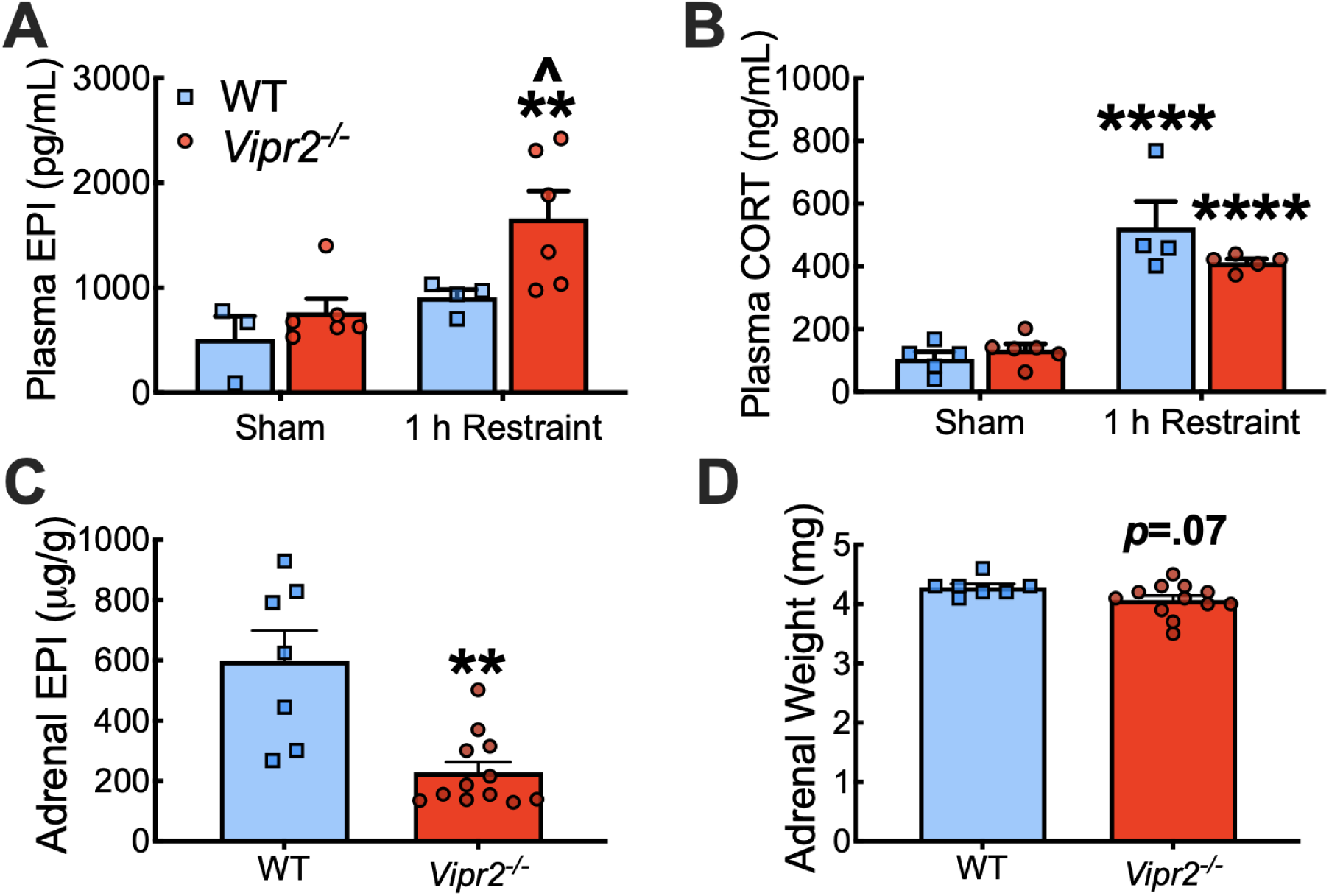
Mild psychogenic stress induced exaggerated epinephrine but not corticosterone secretion in *Vipr2^-/-^*mice. (**A**) Plasma epinephrine measured after 1 h restraint stress was exaggerated in *Vipr2^-/-^* vs WT female mice. (**B**) Plasma corticosterone measured after 1 h restraint stress was elevated in both *Vipr2^-/-^*vs WT. (**C**) Epinephrine content in adrenal glands harvested at necropsy was reduced in *Vipr2^-/-^* vs WT. (**D**) Adrenal weight measured at necropsy displayed an apparent reduction in *Vipr2^-/-^* vs WT. *significantly different vs corresponding sham group in panels A and B or vs WT in panel C, \*\**p*<0.01, \*\*\*\**p*<0.0001. ^significantly different vs WT during restraint, ^*p*<0.05. *n*, 3-6/group in Panel A, 4-6/group in panel B and 7-12/group in panels C and D

For plasma CORT, measured after 1 h restraint stress, there was a main effect of *stress* (F_(1,16)_=91.30, *p*<0.0001) but not *genotype* (*F*_(1,16)_=1.31, ns) or *stress* x *genotype* interaction (*F*_(1,16)_ =3.63, ns; two-way ANOVA). Sidak’s *post-hoc* test for multiple comparisons revealed a significant elevation in plasma CORT between stress and sham groups for *Vipr2^-/-^* (*p*<0.0001) and WT genotypes (*p<*0.0001) (**Fig 6B**). There was no significant difference between CORT values across groups during rest (*p*=0.82) nor during stress (*p*=0.10).

### Vipr2 gene-deletion altered expression of hypothalamic VIP and brainstem Th

We used PCR analysis to assess mRNA expression of gene markers for PACAP, VIP, SNS, and HPA systems in hypothalamus, medulla oblongata and adrenal gland. The results reveal a significant downregulation of hypothalamic *Vip* (Student’s t-test: *t*_(17)_=3.8, adjusted *p* value <0.05) and an apparent upregulation of hypothalamic *Adcyap1* (Welch’s t-test: *t*_(14.4)_ =2.09, adjusted *p* value=0.08) in *Vipr2^-/-^* compared to WT (**Fig. 7**). There was no transcript alteration in hypothalamic *Crh,* a marker of HPA axis activity, in *Vipr2^-/-^* vs WT. In medulla *Vipr2^-/-^*mice showed a significant upregulation in *Th* (Welch’s t-test: *t*_(12.6)_=2.5, adjusted *p* value<0.05) and an apparent upregulation of *Vglut2* (Student’s t-test: *t*_(14)_=2.2, adjusted *p* value=0.09). In adrenal, *Vipr2^-/-^* mice showed an apparent downregulation in *Vip* (Student’s t-test: *t*_(16)_=2.6, adjusted *p* value=0.08). Adrenal copy number for mRNA levels indicated no significant group differences for any GOI studied (*Adcyap1*, *Adcyap1r1, Vip, Th and Actb).* Because adrenal weight may have affected our comparisons, **Table 3** also illustrates the adrenal gene copy number normalized to adrenal weight. However, no significant group differences were found. Correlation and regression analysis between stress peak glycemia vs hypothalamus *Vip* yielded opposite relationships in mutants (positive and apparent) and WT (negative and significant) (*r*=0.45, *p*=0.07 R^2^=0.20, *p*=0.14 for *Vipr2^-/-^; r*=-0.70, *p*=0.04, R^2^=0.49, *p*=0.08 for WT) (**Fig. 8E)**. However, only WT showed a good linear fit. No significant correlation was measured between stress plasma epinephrine vs hypothalamic *Vip* expression or between stress glycemia vs medullary *Th* expression or between plasma epinephrine vs medullary *Th* expression.

**Fig. 7.**
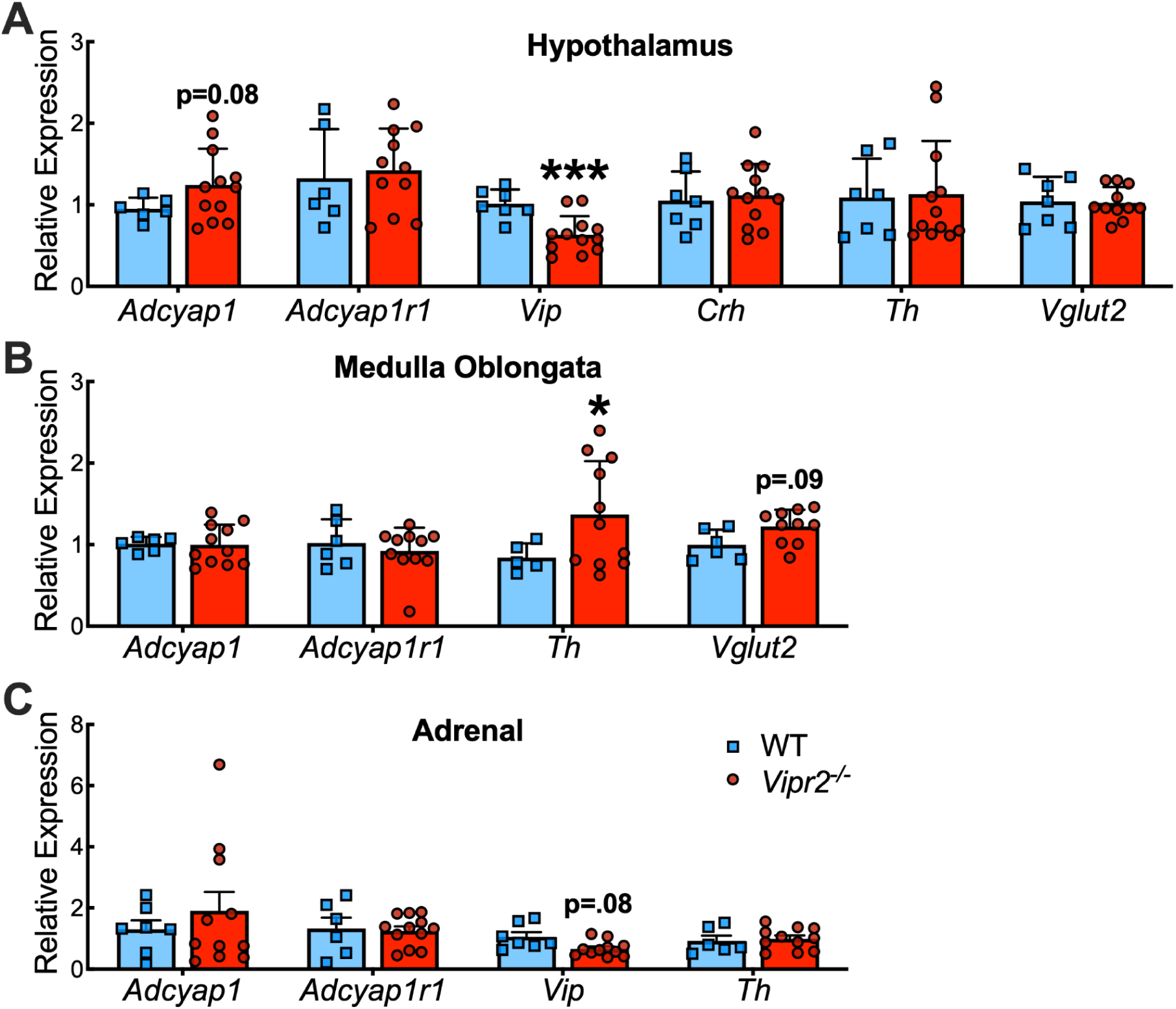
*Vipr2* gene*-*deletion altered expression of gene markers for PACAP, VIP and brainstem sympathetic nervous systems. RT-qPCR analysis illustrates the mean gene expression relative to the reference gene *ActB* of WT vs *Vipr2^-/-^* females and in hypothalamus (**A**), medulla oblongata (**B**) and adrenal gland (**C**). *significantly different vs WT, \**p*<0.05, \*\*\**p*<0.001. *n*, 5-12/group. *Adcyap1*, adenylate cyclase activating polypeptide 1; *Adcyap1r1*, adenylate cyclase activating polypeptide 1 receptor 1; *Crh*, corticotropin-releasing hormone; *Th*, tyrosine hydroxylase; *Vglut2 (Slc17a6)*, vesicular-glutamate transporter 2; *Vip*, vasoactive-intestinal polypeptide

**Fig 8.**
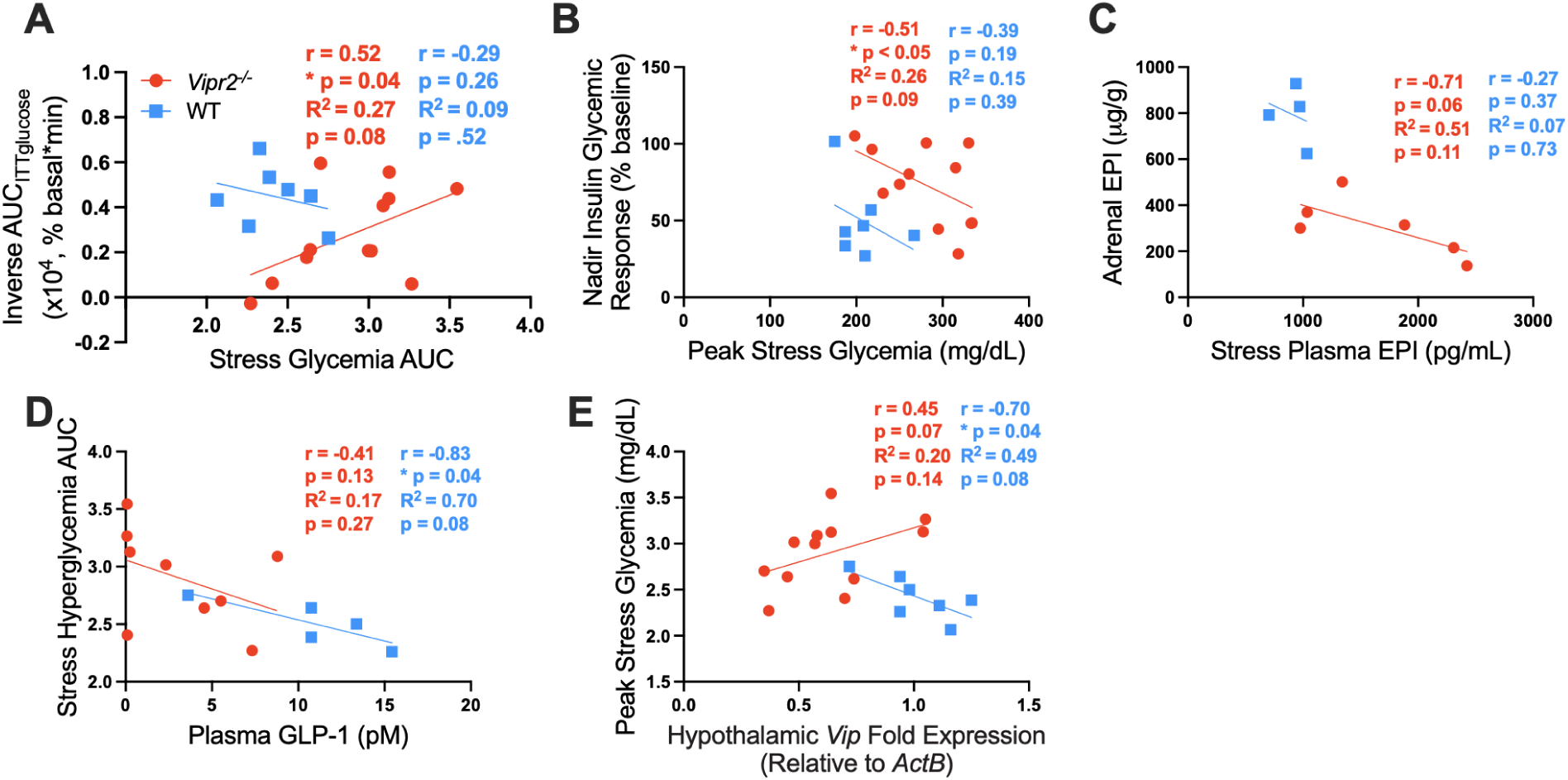
Correlation analysis between selected assay outcomes. Pearson correlation coefficient (r), coefficient of determination (R^2^) with corresponding p-values indicate the strength of the relationship between two variables and the prediction of the value of one continuous variable on another, respectively. (**A**) Inverse AUC_ITTglucose_(% basal) vs stress glycemia AUC. (**B**) Insulin glycemia nadir (% baseline) vs peak stress glycemia. (**C**) Adrenal epinephrine vs stress plasma epinephrine. (**D**) Stress hyperglycemia AUC vs plasma GLP-1. (**E**) Peak stress glycemia vs hypothalamic *Vip* expression. *indicates a significant linear fit (*p*<0.05). EPI, epinephrine

**Table 3.** PCR transcripts in adrenal gland. Summary of changes in adrenal gene expression for WT and *Vipr2*^-/-^ female mice. Changes in gene expression, corrected for adrenal weight, did not indicate any group differences.

|  | Wild Type |  |  | *Vipr2*<sup>-/-</sup> |  |  |
| --- | --- | --- | --- | --- | --- | --- |
|  | Adrenal Weights (mg) | Copies per RNA Input (μg)/Total RNA isolated (μg) x10<sup>6</sup> | Copy Number (μg)/Total RNA Isolated (μg)/Right Adrenal (mg) x10<sup>5</sup> | Adrenal Weights (mg) | Copies per RNA Input (μg)/Total RNA isolated (μg) x10<sup>6</sup> | Copy Number (μg)/Total RNA Isolated (μg)/Right Adrenal (mg) x10<sup>5</sup> |
| *Adcyap1r1* | 4.29 ± 0.06 | 1.8 ± 0.56 | 4.2 ± 1.3 | 4.07 ± 0.08 | 1.8 ± 0.20 | 4.4 ± 0.54 |
| *Adcyap1* | 4.23 ± 0.03 | 1.5 ± 0.34 | 3.5 ± 0.80 | 4.09 ± 0.09 | 1.2 ± 0.28 | 3.1 ± 0.74 |
| *β-Actin* | 4.29 ± 0.06 | 1.6 ± 0.17 | 3.8 ± 0.43 | 4.07 ± 0.08 | 1.4 ± 0.11 | 3.4 ± 0.30 |
| *Th* | 4.28 ± 0.07 | 2.1 ± 0.33 | 4.9 ± 0.80 | 4.10 ± 0.08 | 2.2 ± 0.28 | 5.5 ± 0.76 |
| *Vip* | 4.29 ± 0.06 | 0.16 ± 0.03 | 0.37 ± 0.06 | 4.10 ± 0.08 | 0.10 ± 0.01 | 0.27 ± 0.03 |

## Discussion

PACAP and VIP regulate energy metabolism through central and peripheral actions, which may involve the sympathoadrenal as well as the hypothalamo-pituitary-adrenal axes [7,14,34,37,65,66]. However, the role of PACAP and VIP receptors during physiological challenges to glucose homeostasis is unclear. Our studies performed on VPAC2R gene-deleted female mice have identified a diabetogenic phenotype that includes fed and fasting hyperglycemia and impaired insulin-induced hypoglycemia compared to WT mice. Additionally, *Vipr2^-/-^* female mice displayed exaggerated stress-induced hyperglycemia and plasma epinephrine levels, suggesting hyperactivity of the sympathoadrenal (SA) system in these mice. In support of this, *Vipr2^-/-^* mice displayed reduced adrenal epinephrine and upregulation of *Th* in medulla oblongata (and downregulation of *Vip* in hypothalamus) at rest. In addition, VPAC2R gene deletion produced low plasma GLP-1 and elevated plasma insulin levels. In combination, these data are consistent with a physiological role of VPAC2R in glucose regulation, especially during metabolic and psychogenic stress, likely involving the participation of sympathoadrenal activity and/or metabolic hormones.

We examined the effect of genotype on the fasting glucose response to insulin challenge. A previous study demonstrated that VPAC2R gene-deleted females displayed normal responses, while males showed improved insulin sensitivity after an exogenous insulin challenge (4 h AM fast, 0.75 IU Humulin IP) [67]. Using a similar paradigm, we also observed no effect of *Vipr2* gene deletion on insulin sensitivity in female mice. However, when fast conditions were made more intense (8h ON, 0.25 IU Humulin), female *Vipr2^-/-^* mice displayed impaired insulin hypoglycemia in spite of normal glucose uptake initially after insulin challenge (K_ITT_) compared to WT. Given our GTT results, this is not likely due to less glucose uptake by skeletal muscle, the primary route of glucose disposal [68]. Instead, impaired insulin hypoglycemia may result from dramatically low plasma levels of the incretin GLP-1 in the mutant mice leading to less GLP-1-mediated suppression of endogenous glucose production. Also, reduction of circulating GLP-1 may have hindered its facilitation of glucose disposal [69]. Given the ability of GLP-1 agonists to control fasting glycemia and glucose tolerance, understanding the role of VPAC2R in regulating GLP-1 may aid efforts to discover new therapeutic targets for type 2 diabetes (T2D) [70,71]. Plasma GLP-1 levels were not correlated to impaired insulin hypoglycemia nor to elevated plasma insulin levels found in *Vipr2^-/-^* mice, suggesting unrelated regulation of these parameters. Similarly, no significant relationship was detected between plasma GLP-1 and insulin glycemia response, stress glycemia or plasma epinephrine in *Vipr2^-/-^* mice. Little is known about the role of VPAC2R in GLP-1 hormone regulation, although VPAC1R and VIP-deficient mouse phenotypes and PACAP receptor antagonism are characterized by increased fasting and postprandial GLP-1 levels [7,72,73]. PAC1R- and VIP-deficient fed or fasted mice also exhibit hyperinsulinemia [14,74]. Taken together, these data suggest that the preserved VPAC1 receptors in *Vipr2^-/-^* mice likely mediate the observed inhibitory effects on GLP-1 secretion, triggered by VIP.

It is unlikely that hyperglycemia during insulin challenge was a direct result of loss of VPAC2R function, since activation of VPAC2 receptors does not inhibit hepatic glycogenolysis and glucagon secretion [28]. In fact, because of their inability in this process, in conjunction with their insulinotropic action, VPAC2 receptors are studied for their potential as a novel target for the treatment of T2D [28,30]. More likely is the possibility that, in the absence of VPAC2R in our mutant mice, PACAP- or VIP-mediated activation of the preserved VPAC1 and/or PAC1 receptors, respectively, can promote glucose output via a direct hepatic glycogenolytic effects and/or indirectly by stimulating pancreatic secretion of the glycogenolytic hormone glucagon to counter insulin action [26,27,75]. Indeed, deletion of the PAC1R gene produces impaired but viable insulin-induced hypoglycemia and impaired secretion of glucagon [75]. Taken together, these data support the view that PACAP and/or VIP, by activating either PAC1 or VPAC receptors, in addition to their insulinotropic action, regulate glucose homeostasis in a complex manner that remains to be fully elucidated.

Another counterregulatory response to insulin hypoglycemia is direct sympathetic and/or indirect sympathoadrenal activation of hepatic glucose output. Like glucagon, epinephrine counters glucose uptake by inhibiting insulin secretion, but also by promoting the liberation of glucose from liver via glycogenolysis and gluconeogenesis [76]. Indeed, impaired long-term secretion of epinephrine and depletion of its adrenomedullary stores has been implicated in the lethal hypoglycemic responses to insulin observed in PACAP-deficient mice [33]. Under a single prolonged (90 min) restraint, our *Vipr2^-/-^* mice displayed exaggerated hyperglycemia as compared to WT mice. The gene-deleted mice also displayed a markedly elevated plasma epinephrine in response to stress. These results suggest that defunctionalization of VPAC2R may result in hyperglycemia via PACAP/VIP-mediated sympathetic activation [33,34,77]. Our results prompt us to speculate that exaggerated stress hyperglycemia in *Vipr2^-/-^* mice is either due to removal of potentially inhibitory actions of VPAC2R on sympathetic activation of the liver and/or adrenal and/or due to the unopposed stimulatory actions of VPAC1R and PAC1R on the liver [26,27,75]. In any case, in the mutants, we found an apparent negative association between adrenal epinephrine vs plasma epinephrine, suggesting that circulating epinephrine responses to psychogenic stress were not compensated for by enhanced adrenal epinephrine biosynthesis. Furthermore, exaggerated stress glycemia in *Vipr2^-/-^* was not likely due to an overly active hypothalamo-pituitary-adrenal (HPA) axis, since both genotypes displayed similar elevated stress levels of CORT, which contributes to hepatic glucose output and is regulated by PACAP and VIP [66].

PACAP and VIP are co-transmitters with acetylcholine at the adrenomedullary synapse, where sympathetic regulation of epinephrine occurs in response to stress, secondary to the induction of *Th* [78]. Using *in vivo* restraint stress, Stroth and colleagues (2013) showed that PACAP gene deletion significantly *reduces*, but does not abolish, the upregulation of adrenal *Th* and *Pnmt*, suggesting that both PACAP and VIP contribute to catecholamine biosynthesis and secretion [35]. Given that application of VIP and PACAP to *in vitro* adrenal slices *stimulates* catecholamine release via the activation of all three VIP/PACAP receptor subtypes [79], our findings that gene deletion of *Vipr2* promoted epinephrine release and exaggerated stress-induced hyperglycemia may represent an inhibitory central action by VPAC2R and/or upregulated stimulatory activity of the preserved PAC1 receptors [80]. In support of the former, Inglott and colleagues (2012) reported that activation of VPAC receptors, in contrast to that of PAC1 receptors, *decreased* splanchnic sympathetic nerve activity when activated by intrathecal VIP [81]. It should be noted that baseline levels of adrenal mRNA transcripts for *Adcyap1r1* and *Adcyap1* were not significantly altered in *Vipr2^-/-^* mice, ruling out compensatory changes in PACAP or PAC1 receptor in upregulating adrenal epinephrine and hyperglycemia. Additional mechanistic experiments will need to be undertaken to decipher the role of VPAC2R in hyperglycemia responses to psychogenic stress.

Centrally, VPAC2R may have a differential contribution to hepatic glucose output and glucose disposal. For example, VPAC2 receptors in the RVLM of medulla mediate sympathoexcitation induced by PACAP [41]. Another study by Yi and colleagues (2010), demonstrated the participation of central VPAC2R on pre-autonomic neurons in the paraventricular nucleus (PVN) of the hypothalamus in the control sympathetic stimulation of hepatic glucose production [37]. PACAP-immunoreactive pre-autonomic neurons in the PVN tightly regulate peripheral glucose homeostasis via projections to the brainstem. These brainstem neurons include the adrenergic C1 group pre-sympathetic neurons in medulla [52,53]. Our PCR results indicate transcript modifications in *Vipr2^-/-^* mice that are consistent with sympathoadrenal activation: upregulation of medulla *Th* and downregulation of hypothalamic *Vip.* These central gene alterations may help explain the elevated hyperglycemic responses seen during insulin challenge and psychogenic stress in *Vipr2^-/-^* mice. *Vipr2^-/-^* mice also displayed baseline hyperglycemia in fed and fasted states (8h ON fast), likely due to VPAC1R-mediated hepatic glucose production [26–28] with additional contribution of an enhanced basal metabolic rate [67]. Correlation and regression analyses of peak stress glycemia vs hypothalamic *Vip* expression suggested a negative relationship. Interestingly, this occurred in WT only, suggesting that other factors may influence this relationship in *Vipr2^-/-^*.

The combined results of the insulin glycemia and stress-induced glycemia assays provide a more complete phenotype of *Vipr2^-/-^* mice. To determine if these phenotypes in *Vipr2^-/-^* mice are related, we carried out correlation and regression analysis which revealed an apparent and unexpected *negative* predictive relationship of insulin glycemia (nadir) vs stress glycemia (peak), indicating that *exaggerated* stress hyperglycemia is related to *efficient* insulin hypoglycemia. Using the inverse AUC values for insulin glycemia an apparent *positive* relationship was found with stress glycemia (AUC), suggesting a similar interpretation. It is unclear what mechanisms underlie this relationship in gene-deleted mice, which show abnormal responses in both outcomes relative to WT. Nevertheless, our findings suggest that VPAC2R contributes to physiological responses to metabolic and psychogenic challenges. Our results support previous studies emphasizing that VPAC2R agonists may be leveraged as attractive drug candidates for the treatment of diabetes [21,23,28,30,82] and stress-related psychopathologies [83–85].

### Strengths and Limitations to the study

The strength of the current study is a more complete characterization of the physiological, and homeostatic effects of VPAC2R gene deletion using a null murine model. Our findings indicate participation of VPAC2R in the regulation of glucose homeostasis under metabolic and psychogenic stress. A limited number of mice were used and, therefore, all mice were required for all assays. To minimize potential confounds, we allowed 1-3 weeks between assays. All mice received restraint stress to measure hyperglycemia responses. However, only a subset within each genotype group received stress in the following assay measuring plasma epinephrine. Importantly, correlation/regression analysis conducted on the values of consecutive assays did not indicate a predictive relationship on outcomes in both genotype groups arguing against the possibility of procedurally biased outcomes.

Although we did not measure feeding patterns in our mutant mice, their reported circadian deficit should not have impacted the timing of feeding and of glycemia values in fed and fasted states since mice were housed under a steady light:dark cycle. Harmar and colleagues (2002) showed that upon exposure to continuous dim red light VPAC2R mutant males failed to express circadian clock genes and exhibited grossly disrupted rhythms of activity [16]. However, there was little disruption in the presence of a normal 12 hr light:dark cycle, i.e., a shorter lag in activity onset when responding to the dark periods and similar periods of rhythmicity.

Another weakness of the current study is its reliance on a transgenic animal model, although it provides an alternative to pharmacological manipulation of PACAP/VIP receptors with only moderately selective drugs. Using the transgenic approach we were unable to evaluate VPAC2R-mediated effects due to the potential binding of either PACAP or VIP, for which the receptor has equal affinity [86]. In addition, VPAC2R gene deletion produced unexpected changes in central PACAP/VIP systems gene expression. i.e., downregulation of *Vip* in hypothalamus (significant) and adrenal (apparent) and upregulation of *Adcyap1* in hypothalamus (apparent), that may have contributed to phenotypic effects observed.

*Vipr2^-/-^* female mice displayed impaired insulin hypoglycemia and exaggerated stress hyperglycemia. The potential relationship between these two phenotypes is unclear and requires further study. Correlation/regression analysis indicated a positive apparent relationship between insulin glycemia responses (inverse AUC) and stress glycemia only in *Vipr2^-/-^*mice, suggesting that abnormally *elevated* stress hyperglycemia was related to *efficient* insulin hypoglycemia. It is unclear what mechanisms underlie this relationship in gene-deleted mice, which show abnormal responses in both outcomes relative to WT. Furthermore, these abnormal glycemia responses did not associate significantly with altered sympathoadrenal control of plasma epinephrine or plasma GLP-1 levels. Also, *Vip2r^-/-^*mice did not show the apparent negative association between hypothalamic *Vip* and stress glycemia nor between stress glycemia and plasma GLP-1 observed in WT mice. Our findings indicate a physiological role of VPAC2R in glucose homeostasis under metabolic and psychogenic stress.

## Abbreviations

*Actb*: beta actin
*Adcyap1*: Pituitary adenylate cyclase-activating polypeptide gene
*Adcyap1r1*: Pituitary adenylate cyclase-activating polypeptide receptor 1 gene
ANOVA: analysis of variance
AUC: area under curve
CORT: corticosterone
Cq: cycle quantification
*Crh*: corticotropin-releasing hormone gene
CNS: central nervous system
ELISA: enzyme-linked immunosorbent assay
FBG: fasting blood glucose
GLP-1: glucagon-like peptide-1
GPCR: G-protein-coupled receptor
GTT: glucose tolerance test
HPA: hypothalamic-pituitary-adrenal
ITT: insulin tolerance test
IS: immobilization stress
ON: overnight
PACAP: pituitary adenylate cyclase-activating polypeptide
PAC1R: Pituitary adenylate cyclase-activating polypeptide type I receptor
RT: reverse transcriptase
SA: sympathoadrenal
SNS: sympathetic nervous system
*Th*: tyrosine hydroxylase gene
T2D: type II diabetes
THI: trihydroxyindole
*Vglut2*: vesicular glutamate transporter 2 gene (*Slc17a6*)
VIP: vasoactive-intestinal peptide
*Vip*: vasoactive-intestinal peptide gene
*Vipr2*: vasoactive-intestinal peptide receptor 2 gene
VPAC1R: vasoactive-intestinal peptide receptor 1
VPAC2R: vasoactive-intestinal peptide receptor 2
WT: wild type

## Acknowledgements

We acknowledge Dr. J. Waschek, UCLA, for the gift of WT and *Vipr2^-/-^* mice. We thank H. Clark, UCR Genomics Core, for technical assistance with Bioanalyzer. We thank Dr. K. Xu, UC Riverside, for help with statistical analysis. We acknowledge fellowship support from NSF GRFP fellowship (MCV), STEM-Hispanic Serving Institutions (HSI) Department of Education Award and University of California UC-HSI Doctoral Diversity Initiative (UC-HSI DDI) President’s Pre-Professoriate Fellowship (E.V.K.). Figure 1 was created with Biorender.com.

## Declarations

## Availability of Data

The data that support the findings of this study are available from the corresponding author upon reasonable request.

## Disclaimer

The opinions and assertions expressed herein are those of the author(s) and do not necessarily reflect the official policy or position of the Uniformed Services University or the Department of Defense.

J.M.K. is now a 2nd Lieutenant at the Uniformed Services University, Department of Defense. Her work was performed at the University of California, Riverside before becoming a military officer. However, we want to emphasize that the opinions and assertions expressed herein are those of the authors and do not necessarily reflect the official policy or position of the Uniformed Services University or the Department of Defense.

## Ethics approval

Care and treatment of animals was performed in compliance with NIH guidelines and approved by the University of California, Riverside Institutional Animal Care and Use Committee (AUP# 20170026 and 20200018).

## CRediT authorship contribution statement

**Conceptualization**: EVK, MCC; **Data Curation**: EVK, AEB, MED; **Formal Analysis**: EVK, AEB, MED, BDC, MCC; **Funding acquisition**: EVK, MCV, KAS, MCC; **Investigation**: EVK, AEB, MED, BDC,KB, JMK; **Methodology**: EVK, AEB, MED, MCV, KAS, MCC; **Project administration**: EVK, MCC; **Software**: EVK; **Supervision**: EVK; MCC; **Validation**: EVK, MED, MCV, KAS, MCC; **Visualization**: EVK, MCC; **Writing - original draft**: EVK, AEB, MED, MCC; **Writing - review and editing**: EVK, AEB, MED, MCC.

## Consent to participate

Not applicable.

## Consent for publication

All authors reviewed and approved the final manuscript.

